# Niclosamide Inhalation Powder Made by Thin-Film Freezing: Pharmacokinetic and Toxicology Studies in Rats and Hamsters

**DOI:** 10.1101/2021.01.26.428293

**Authors:** Miguel O. Jara, Zachary N. Warnken, Sawittree Sahakijpijarn, Chaeho Moon, Esther Y. Maier, Dale J. Christensen, John J. Koleng, Jay I. Peters, Sarah D. Hackman, Robert O. Williams

## Abstract

In this work, we have developed and tested *in vivo* a dry powder form of niclosamide made by thin-film freezing (TFF) and administered it by inhalation to rats and hamsters. The niclosamide dry powder, suitable for inhalation, is being developed as a therapeutic agent against COVID-19 infection. Niclosamide, a poorly water-soluble drug, is an interesting drug candidate because it was approved over 60 years ago for use as an anthelmintic medication, but recent studies demonstrated its potential as a broad-spectrum antiviral with a specific pharmacological effect against SARS-CoV-2 infection. In the past, clinical trials for other indications were terminated prior to completion due to low and highly variable oral bioavailability. In order to quickly address the current pandemic, targeting niclosamide directly to the lungs is rational to address the COVID-19 main clinical complications. Thin-film freezing technology was used to develop a niclosamide inhalation powder composition that exhibited acceptable aerosol performance with a fine particle fraction (FPF) of 86.0% and a mass median aerodynamic diameter (MMAD) and geometric standard deviation (GSD) of 1.11 μm and 2.84, respectively. This formulation not only proved to be safe after an acute three-day, multi-dose pharmacokinetic study in rats as evidenced by histopathology analysis, but also was able to achieve lung concentrations above the required IC_50_ and IC_90_ levels for at least 24 h after a single administration in a Syrian hamster model. To conclude, we successfully developed a niclosamide dry powder inhalation formulation by thin-film freezing for further scale-up and clinical testing against the COVID-19 infection. This approach overcomes niclosamide’s limitation of poor oral bioavailability by targeting the drug directly to the primary site of infection, the lungs.

## Introduction

The Coronavirus Disease 2019 (COVID-19) is a pandemic caused by the coronavirus (CoV), also known as Severe Acute Respiratory Syndrome Coronavirus 2 (SARS-CoV-2). SARS-CoV-2 is related to other CoVs that have caused other pandemics, including Severe Acute Respiratory Syndrome (SARS) and the Middle East Respiratory Syndrome (MERS) in 2002-2003 and 2012, respectively [1,2]. Currently, the COVID-19 pandemic has unfortunately killed more people than either of the other before-mentioned pandemics combined [1,3,4].

In 2004, Wu *et al*. reported that the FDA-approved drug niclosamide inhibits the viral replication of SARS-CoV [5]. More recently, in 2019, niclosamide was shown to inhibit the viral replication of MERS-CoV by more than a 1000-fold by interfering in pathways related to the proteasome and autophagy mechanisms [6,7]. Now, in 2020, during the COVID-19 pandemic, niclosamide has shown very promising effectiveness against the viral replication of SARS-CoV-2 with a reported IC_50_ of 0.28 μM (~91.56 ng/mL) by similar reported mechanisms of action as that reported for SARS-CoV, and the mechanisms of anti-viral activity may include potentially direct intermolecular interactions between the drug and SARS-CoV-2 proteins [7–10]. Based on the data reported by Joen et al., Arshad *et al*. (2020) estimated an IC_90_ of 153.7 ng/mL for niclosamide [11]. This evidence supports niclosamide as a candidate for repurposing as an antiviral for the current pandemic.

Unfortunately, niclosamide is a poorly water-soluble molecule with limited oral bioavailability, thus limiting its ability to be repurposed for other clinical indications. The clearest example of this limitation is clinical trial NCT02532114 that was terminated prematurely due to the low and variable bioavailability of niclosamide, reportedly only reaching C_max_ values between 35.7-182 ng/mL when administered orally at 500 mg three-times-daily. However, SARS-CoV-2 mainly affects the pulmonary airways; the virus has a tropism towards alveolar macrophages and pneumocytes (type I and II) [12]. Here the virus leads to breathing difficulties, acute respiratory distress syndrome, pneumonia, among other complications [13,14]. A way to overcome niclosamide’s severe bioavailability limitation and deliver the drug directly to the lungs is by the development of niclosamide as an inhaled product. Niclosamide has been reported to be safe for inhalation after 24 h of administration [15]. However, to the best of our knowledge, a pharmacokinetic evaluation of inhaled niclosamide has not previously been reported.

Thin-film freezing (TFF) is a technology used for particle engineering, usually aiming to develop powders for inhalation for use with dry powder inhalers (DPI). The TFF-processed powders generally are low-density, brittle, and porous matrices with excellent aerosol performance [16]. The TFF technology generates these powders by fast supercooling of drug-carrier solutions (sometimes suspensions) [17]. The frozen material then is lyophilized to remove the employed solvents (e.g., acetonitrile, 1,4 dioxane, tert-butanol or water). TFF technology has proven to be useful in generating powders for delivery by DPI, and currently, there are two formulations in phase I clinical trials (i.e., voriconazole and tacrolimus) using this established technology [18–21]. For these reasons, the development of an inhaled niclosamide powder by TFF is hypothesized to offer a new therapeutic option and route of administration in the fight against COVID-19.

In this work, we aim to develop a formulation to reach the lungs and sustain niclosamide concentrations, which is difficult to achieve by other routes of administration due to niclosamide’s poor-water solubility and high first-pass metabolism [22,23]. For this purpose, the niclosamide inhalation powder requires aerodynamic diameters between 1 to 5 μm, necessary for lung delivery where it can exert its pharmacological effect [24].

The goals of the research are summarized as follows: (1) develop a dry powder of niclosamide for inhalation by TFF manufacturing as a potential therapeutic for COVID-19; (2) test its *in vitro* aerodynamic properties; 3) characterize its solid-state properties; and (4) evaluate its pharmacokinetics and lung toxicity *in vivo* using the rat and Syrian golden hamster models.

## Materials and Methods

### Materials

Niclosamide was purchased from Shenzhen Nexconn Pharmatechs LTD. (Shenzhen, China). Tert-butanol (TBA), acetone, acetonitrile, leucine, polysorbate 20, trifluoracetic acid were acquired from Fisher Scientific (Pittsburgh, PA, USA). Pearlitol^®^ PF-mannitol was purchased from Roquette America (Keokuk, IA, USA).

### Preparation of Thin Film Freezing (TFF) composition

In the preparation of the solutions for TFF manufacturing, niclosamide was dissolved in TBA, comprising the organic phase. The hydrophilic excipients, mannitol, and leucine were dissolved in water, which formed the aqueous phase. Then, the aqueous and organic phases were mixed as specified in Table 1. Thereafter, the mixed solution was applied as drops onto a rotating cryogenically cooled drum at cooled to −120 °C. The frozen solids were collected with liquid nitrogen and stored in a −80°C freezer before lyophilization. The lyophilization was performed in a SP VirTis Advantage Pro shelf lyophilizer (SP Industries, Inc., Warminster, PA, USA). The primary drying process was at −40 °C for 20 h, and then, the temperature was linearly increased to 25 °C over 20 h, followed by holding the temperature at 25 °C for 20 h. The pressure was maintained at 100 mTorr during the lyophilization process.

**Table 1.** Composition of the inhalation powders made by thin-film freezing and their corresponding aqueous and organic phases used in the process.

| **Animal model study** | **Composition of inhaled powder made by TFF (%w/w)** | **Organic phase composition (mg/mL)** | **Aqueous phase composition (mg/mL)** | **Organic/Aqueous phase ratio (v/v)** |
| --- | --- | --- | --- | --- |
| Sprague-Dawley Rats | 22% Niclosamide-73% Mannitol-5% Leucine | Niclosamide 1.38 mg/mL | Mannitol 18.25 mg/mL<br>Leucine 1.25 mg/mL | 4/1 |
| Syrian golden hamster | 1.66% Niclosamide-92.04% Mannitol-6.3%<br>Leucine | Niclosamide 0.10 mg/mL | Mannitol 23.00 mg/mL<br>Leucine 1.58 mg/mL | 4/1 |

### Aerodynamic particle size distribution analysis

Five milligrams of niclosamide inhalation powder was loaded into size #3 hydroxypropyl methylcellulose capsules (Vcaps^®^ plus, Capsugel^®^, Lonza, Morristown, NJ, USA). The aerodynamic properties of the powder were evaluated using a Next Generation Impactor (NGI) (MSP Corporation, Shoreview, MN, USA) connected to a High-Capacity Pump (model HCP5, Copley Scientific, Nottingham, UK) and a Critical Flow Controller (model TPK 2000, Copley Scientific, Nottingham, UK). The dry powder inhaler device RS00 (Plastiape^®^, Osnago, Italy) was used for dispersing the powder through the USP induction port with a total flow rate of 58 L/min for 4.1 s per each actuation corresponding to a 4 kPa pressure drop across the device and a total flow volume of 4 L. To avoid particle bounce, a solution of polysorbate 20 in methanol at 1.5% (w/v) was applied and dried onto the NGI collection plates to coat their surface. The pre-separator was not used in this analysis. After dispersal, the powder was extracted from the stages using a mixture of water/acetonitrile (20:80 v/v). Then, the samples were analyzed by HPLC described below. The analysis was conducted three times (n=3).

The NGI results were analyzed using the commercial software Copley Inhaler Testing Data Analysis Software 3.10 (CITDAS) (Copley Scientific, Nottingham, UK). CITDAS provided the calculation for mass median aerodynamic diameter (MMAD), total dose per shot, calculated delivered dose, fine particle dose, fine particle fraction of delivered dose (FPF %, delivered dose), and geometric standard deviation (GSD).

The fine particle fraction of the recovered dose (FPF %, recovered dose) was calculated as the percentage of the total amount of niclosamide with an aerodynamic diameter below 5 μm over the total amount of niclosamide collected. Total dose per actuation is the total amount of niclosamide collected after performing NGI. The calculated delivered dose is the amount of drug collected from the throat (mouthpiece adapter + induction port) to the micro-orifice collector (MOC). It does not consider the niclosamide retained in the capsule and device after actuation. The fine particle dose is the mass of niclosamide with an aerodynamic diameter below 5 μm.

### Scanning Electron Microscopy (SEM)

The TFF powder was placed onto carbon tape and mounted onto aluminum stubs followed by and sputter-coating with platinum/palladium to a thickness of 15 nm (Cressington Scientific Instruments LTd., Watford, UK) before capturing the images using a Zeiss Supra 40 (Carl Zeiss, Oberkochen, Germany).

### Differential scanning calorimetry (DSC)

Analyses were performed using a differential scanning calorimeter model Q20 (TA Instruments, New Castle, DE, USA) operating in ramp DSC mode from 35°C to 240°C with a ramp temperature of 10°C/min and a nitrogen purge of 50 mL/min. Tzero pans and lids were used in the experiment Q20 (TA Instruments, New Castle, DE, USA).

### X-ray powder diffraction (XRD)

The XRD studies were conducted using a Rigaku MiniFlex 600 II (Rigaku Americas, The Woodlands, TX, USA) equipped with primary monochromated radiation (Cu K radiation source, λ = 1.54056 Å). The 2-theta angle was set at 5-40° (0.02° step, 1°/min, 40 kV, 15 mA).

### HPLC analysis

The samples were measured at 331 nm using a Dionex HPLC system (Thermo Fisher Scientific Inc., Sunnyvale, CA, USA) with a ZORBAX SB-C18 column (4.6 x 250 mm, 5 μm) (Agilent, Palo Alto, CA, USA) at 1 mL/min flow rate. An isocratic method was used with a mobile phase consisting of a formic acid aqueous solution at 0.3% (v/v) and acetonitrile mixed in a 40:60 ratio.

### Multi-dose pharmacokinetic study in rats

A three-day pharmacokinetic study was performed in female Sprague-Dawley rats (252.4 – 279.5 g). The animal study was approved by the University of Texas at Austin Institutional Animal Care and Use Committee (IACUC; Protocol Number: AUP-2019-00253). Prior to dosing the niclosamide inhalation powder to the animals, the powder was sieved using a No. 325 sieve (45 μm aperture) to deaggregate larger particles and prevent the insufflator device from clogging inside. A dry powder insufflator (DP-4; Penn-Century Inc., Philadelphia, PA, USA) was weighed empty and filled with niclosamide inhalation powder enough to release 1 mg of the formulation (equivalent to 220 μg of niclosamide) for each administration. Rats were anesthetized with isoflurane (4% induction, 1-5-2.5% maintenance), the metal tube of the DP-4 device was briefly inserted into the trachea, and the powder was puffed into the lung using an air pump (AP-1 model, Penn-Century Inc., Philadelphia, PA, USA). The device was weighed immediately after the procedure to determine the amount of powder released. The animals received the powder once per day for three consecutive days. Twenty-four hours after the final administration, animals were sacrificed, and blood was drawn using a cardiac puncture and transferred into BD microcontainers^®^ (reference number 365985, Becton Dickinson, Franklin Lakes, NJ, USA) and centrifuged for 5 minutes at 10,000 rpm in order to separate the plasma that was immediately frozen until further analysis using HPLC-mass spectrometry (HPLC-MS). The lung was perfused with phosphate buffer solution (PBS), and the right lobes were removed to determine niclosamide content. Prior to removal, the left lobe of the lung was inflated and fixed with 10% buffered formalin. Left lobe, spleen, liver, and kidneys were submerged in 10% buffered formalin for 1-week prior to histopathologic analysis.

### Quantification of niclosamide in rat’s plasma and lung tissue

A plasma calibration curve was prepared by spiking 100 μL of blank plasma with niclosamide standards at concentrations of 0, 5, 10, 50, 100, 500, 1000, and 5000 ng/mL. Samples and calibrators were prepared by adding 10 μL of the internal standard, deuterated phosphatidylethanol 16:0 20:4 (D5-PEth 16:0 20:4), and 1 mL of a 75% MeOH solution. Then, they were then vortexed for 30 minutes, followed by centrifugation at 3200 g for 30 minutes. The supernatant was transferred to clean tubes and dried under a gentle stream of nitrogen. Then, the samples and calibrators were resuspended in 1:1 acetonitrile: 0.1% formic acid water solution. Thereafter, the samples were transferred into HPLC vials, and 10 μL were injected into a Shimadzu SIL 20A HT autosampler, LC-20AD pumps, and an AB Sciex API 4000 Qtrap tandem mass spectrometer with turbo ion spray. The LC analytical column was a C18 Excel 3 ACE (3 x 75 mm, 3 microns) (MacMod, Chadds Ford, PA, USA) and was maintained at 25°C during the measures. Mobile phase A consisted 0.1% formic acid water solution, and mobile phase B consisted of 0.1% formic acid acetonitrile solution. The flow rate was 0.4 mL/min and the gradient was 5% B initially; then, at 1 minute after injection it was ramped to 99% B. From 6 min to 8 min, the mobile phase was maintained at 99% B and at 8.1 minutes was switched immediately back to 5% B, and it was maintained for 2.9 minutes to equilibrate the column before the next injection. The niclosamide transition was detected at 325 Da (precursor ion), and the daughter ion was detected at 171 Da. The internal standard deuterated phosphatidylethanol (D5-Peth) 16:0 20:4 transition was detected at 728 Da (precursor ion), and the daughter ion was detected at 303 Da.

In the case of lung tissue, the calibration curve was prepared by spiking 100 μL of homogenized blank lung tissue with niclosamide standards at concentrations of 0, 5, 10, 50, and 100 ng/mL. Niclosamide was quantified in mouse lung tissue by weighing out the lung and adding a 10x volume of a 75% methanol solution. Then, the samples and calibrators were homogenized, and 100 μL were combined with 10 μL of the internal standard. Then the samples were vortexed for 30 min and followed the same methodology previously described for plasma samples.

### Histopathology

Fixed tissues were embedded in paraffin prior to cutting into 5 μm sections. Sections were stained with hematoxylin solution for 4 min., washed with water for 10 min, followed by staining with eosin solution. The resulting stained tissue sections were analyzed by microscopy.

### Pharmacokinetic study in Syrian hamsters

A single-dose pharmacokinetic study was performed in female Syrian hamsters. The animal study was approved by the Institutional Animal Care and Use Committee (IACUC; Protocol Number: AUP-2019-0235) from The University of Texas at Austin. Two cohorts of 35 female Syrian hamsters (35 – 42 days old and the average weight of 108 ± 8 g) were administered 8.7 or 17.4 mg/Kg of niclosamide inhalation powder each (145 and 290 μg/Kg of pure niclosamide, respectively). The powder was manufactured using the same protocol previously described, but the concentration of niclosamide was reduced in the mixture down to 1.66% (w/w) in order to facilitate dosing appropriate amounts relative to the hamster’s body weights while meeting the minimum fill weight of the DP-4 insufflator (1 mg). Prior to dosing, the niclosamide inhalation powder was sieved using a No. 200 sieve (75 μm aperture) in order to disaggregate larger particles and prevent the device from clogging during use. Similar to the before-mentioned method for rats, a nose cone was used to administer isoflurane at 4% for the induction of anesthesia and subsequently at 2% for its maintenance. The hamsters were placed on their backs and secured at a 45° using silk. The powder was administered intratracheally using the Penn-Century dry powder insufflator™ DP-4 (Penn-Century Inc., Philadelphia, PA, USA). The device was actuated three times to deliver 200 μL of air per actuation. After the administration of niclosamide inhalation powder, blood samples were collected by cardiac puncture at 0.25, 0.5, 1, 2, 4, 8, and 24 h using five hamsters for each time point. The samples were added into BD microcontainers^®^ (reference number 365985, Becton Dickinson, Franklin Lakes, NJ, USA) and centrifuged for 5 minutes at 10,000 rpm in order to separate the plasma that was quantified using HPLC-MS. In the case of lung tissue, the lungs were washed and perfused with PBS, removed, and immediately frozen. Prior to analysis, whole lung tissue was placed into BioStor™ Vials with screw caps (National Scientific Supply, Claremont, CA, USA) with 3.5 g of zirconia/silica beads (BioSpec Products, Bartlesville, OK, USA). The tissue was homogenized at 4,800 rpm for 20 seconds.

### Quantification of niclosamide in hamster’s plasma and lung tissue

The calibrators were prepared by spiking niclosamide standard solutions into blank plasma solutions with a range from 0.1 to 1000 ng/mL. 200 μL of a 100 ng/mL methanolic solution of the internal standard, 13C6 niclosamide (Niclosamide-(2-chloro-4-nitrophenyl-^13^C_6_) hydrate, (Sigma-Aldrich, Saint Louis, MO, USA) was added into 200 μL of plasma samples and calibrators. Then, they were vortexed and centrifuged for 15 min at 12,000 rpm. The supernatant was measured by LC/MS/MS analysis. In the case of lung tissue, the calibrators were prepared by spiking niclosamide standard solutions into blank lung tissue in the range from 0.5 to 10,000 g/mL. 1000 μL of internal standard, 13C6 niclosamide, at 100 ng/mL was added into the weighed lung samples and calibrators. Then, they were vortexed and centrifuged for 15 min at 12,000 rpm. The supernatant was measured by LC/MS/MS analysis. The analysis was performed using Agilent G1367D autosampler, G4220A binary pump, G1316B column compartment, and a G6470A triple quadrupole mass spectrometer. Niclosamide was separated on an Agilent poroshell column 2.1 x 50 mm, 2.7 μm column (Agilent Technologies, Wilmington, DE, USA) using a gradient of 0 to 95% B in 5 min (A= water with 0.025% TFA and B= 95% acetonitrile in water with 0.025% TFA) with a 1-minute hold at the final conditions at a flow rate of 0.35 mL/min. The post run column equilibration was 4 min. The column was held at 40°C for the analysis. The injection volume for MS was 10 μL. The specific details about the ionization can be found in Table S1 (supplemental materials).

### Pharmacokinetic analysis

MATLAB^®^ SimBiology (MathWorks, Natick, MA, USA) software was used to run a noncompartmental analysis of the plasma and lung samples. In this regard, the niclosamide concentrations from the five animals of every sample point were averaged and considered as a single subject to run the NCA as previously reported in studies with small sample sizes [25,26]. In order to compare the previously reported IC_50_ and IC_90_ concentration values to the measured lung concentration in μg/g wet tissue, we assumed that the wet tissue has a density of 1 g/mL.

## Results and Discussion

### Niclosamide inhalation powder has high aerosol performance

The aerosol performance of the niclosamide inhalation powder was determined by NGI, as shown in Table 2. The powder had an FPF (delivered dose) of 86.0 ± 2.7%, which is particularly high when compared with other niclosamide inhalers manufactured by technologies such as spray-drying [15,27]. Remarkably, the inhaled powder achieved an MMAD of about 1 μm and was able to reach the last stage of the cascade impactor (**Error! Reference source not found.**Figure 1). Our main objective was to target niclosamide into the deep lung region with our dry powder formulations in order to cover most of the surface potentially infected by SARS-CoV-2, and this formulation warrants further development.

**Table 2.** Aerosol performance of niclosamide inhalation powder administered to rats.

| Total Dose Per Shot [μg] | Calc Delivered Dose [μg] | Fine Particle Dose [μg] | Fine Particle Fraction [%,<br>delivered dose] | Fine Particle Fraction [%,<br>recovered dose] | MMAD [μm] | GSD |
| --- | --- | --- | --- | --- | --- | --- |
| 709.6<br>(41.1) | 633.2 (53.5) | 543.5 (31.0) | 86.0 (2.7) | 76.6 (1.4) | 1.11 (0.07) | 2.84 (0.22) |

**Figure 1.**
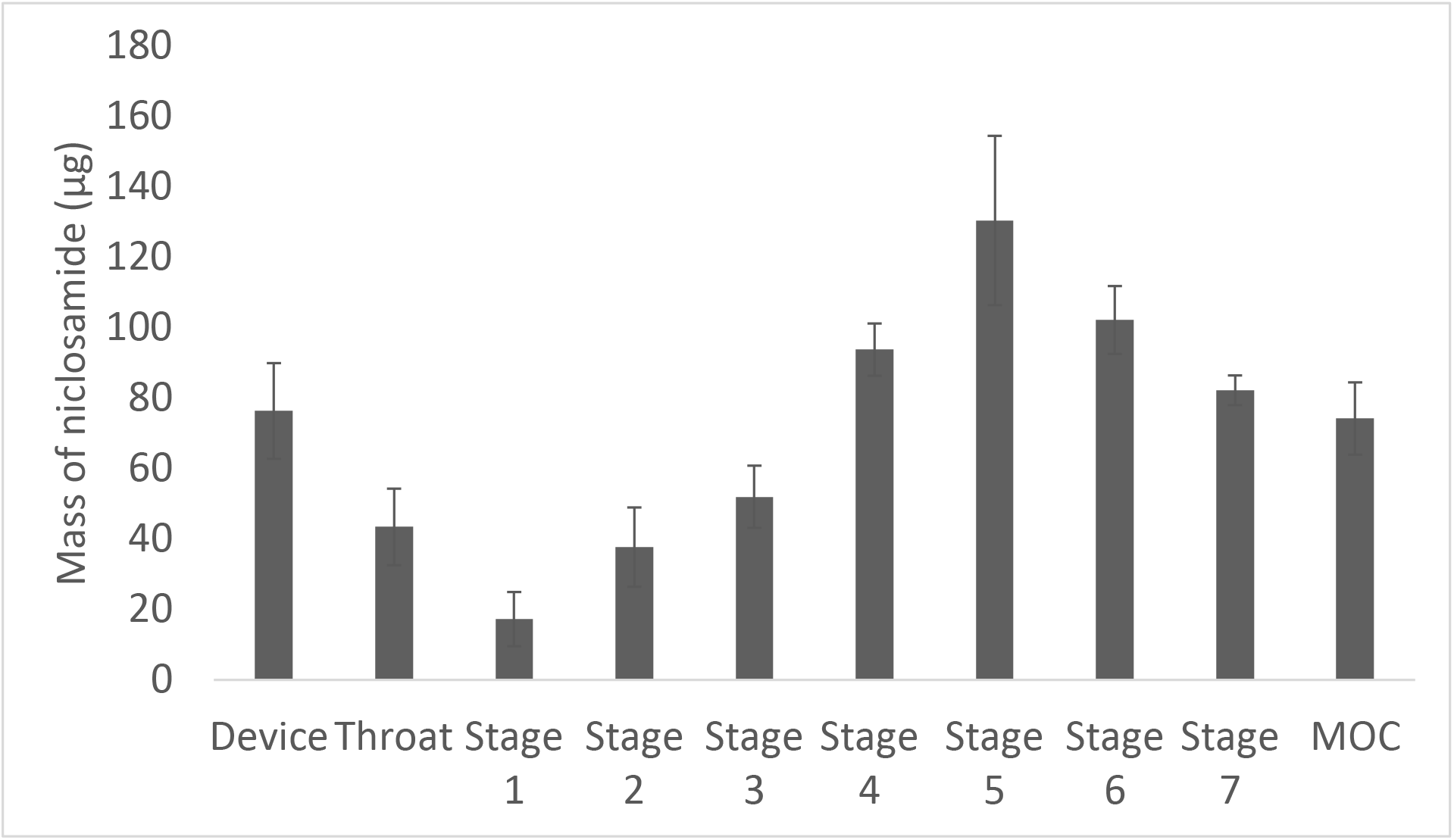
Aerodynamic particle size distribution of niclosamide inhalation powder (n=3) made by thin-film freezing. The device bar indicates the remaining mass of niclosamide in the capsule and device. In the case of the throat bar, it indicates the remaining mass of niclosamide in the mouthpiece adapter and induction port.

### Solid-state characterization of niclosamide inhalation powder shows that the TFF process provides nanonization of drug

The niclosamide inhalation powder was characterized using DSC and XRD (Figure 2). In general, it was not possible to detect the diffraction peak or thermal events from leucine because it was present in low amounts (< 5%) [28]. In the case of mannitol, the TFF process generated a mixture of polymorphs in which patterns of δ-mannitol and β-mannitol were observed [29]. The TFF-mannitol was used as reference material, as shown in Figure 2B. In the DSC, a transition was observed between 110-140°C, which was previously reported by Peters *et al*. (2016) and Burger *et al*. (2000) [30,31]. This endothermic transition is related to the conversion of δ-mannitol into endothermic α-, β or eutectic combinations of both polymorphs of mannitol, followed by melting at ~172 °C. It is important to mention that depending on the freezing/drying conditions and the presence of other excipients, different polymorphs of mannitol and its mixtures can be generated [32,33].

**Figure 2.**
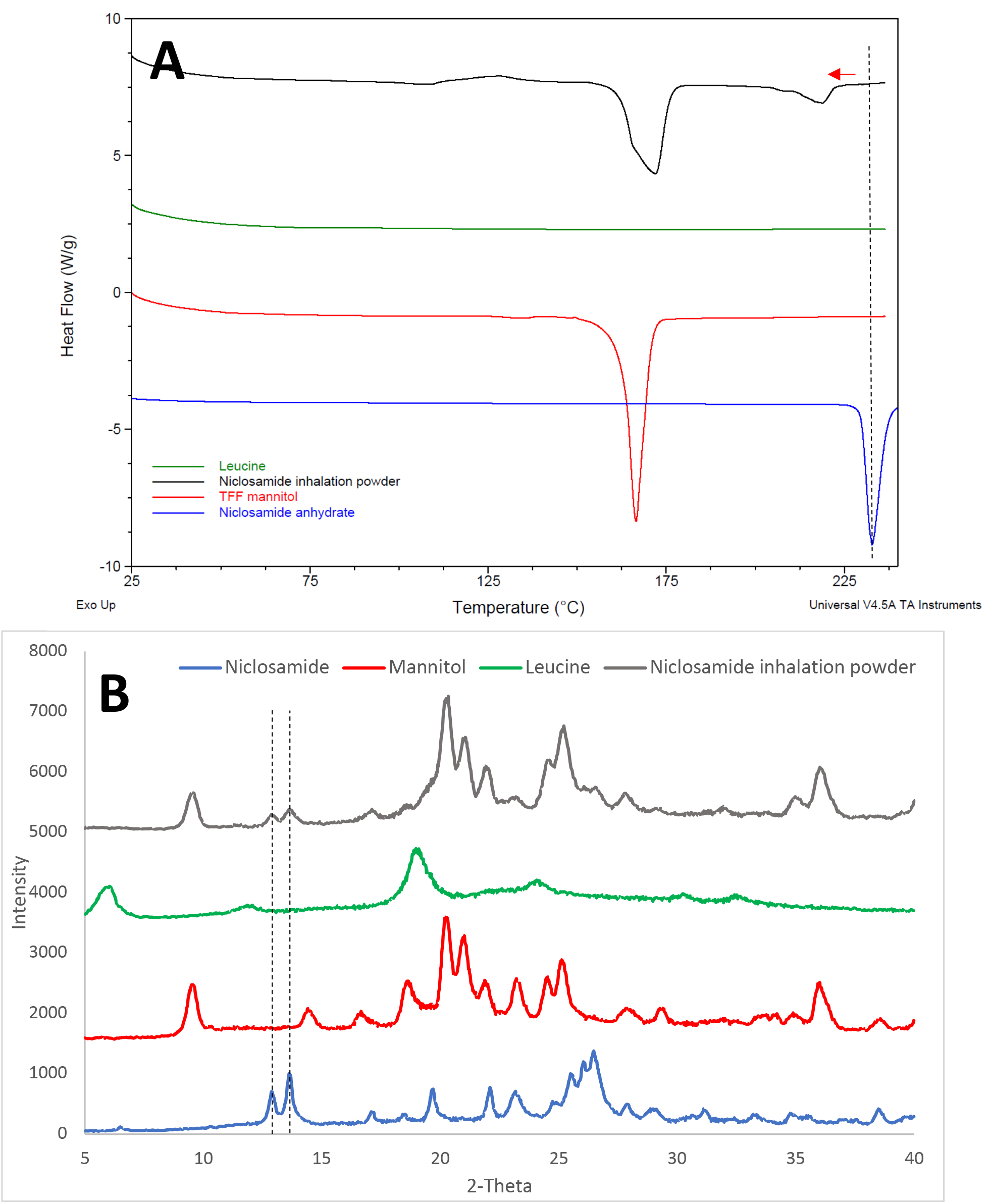
(A) DSC thermogram and (B) XRD diffractogram of niclosamide inhalation powder and its separated constituents. In the thermogram (see, 2A), the dotted line and red arrow indicate the peak shift of niclosamide melting point. In the case of the diffractogram (see, 2B), the dotted lines indicate characteristic peaks of niclosamide anhydrate.

In the case of niclosamide, there was a shift in its melting peak from 230°C to ~218°C. We attributed this behavior to the nanonization of niclosamide crystals by TFF processing supported by the scanning electron microscopy image (Figure 3)[17]. It is reported that nanomaterials exhibit changes in their thermograms, usually manifested as a shift of their melting point to lower temperatures, peak broadening, and even a reduction of their melting enthalpy [34].

**Figure 3.**
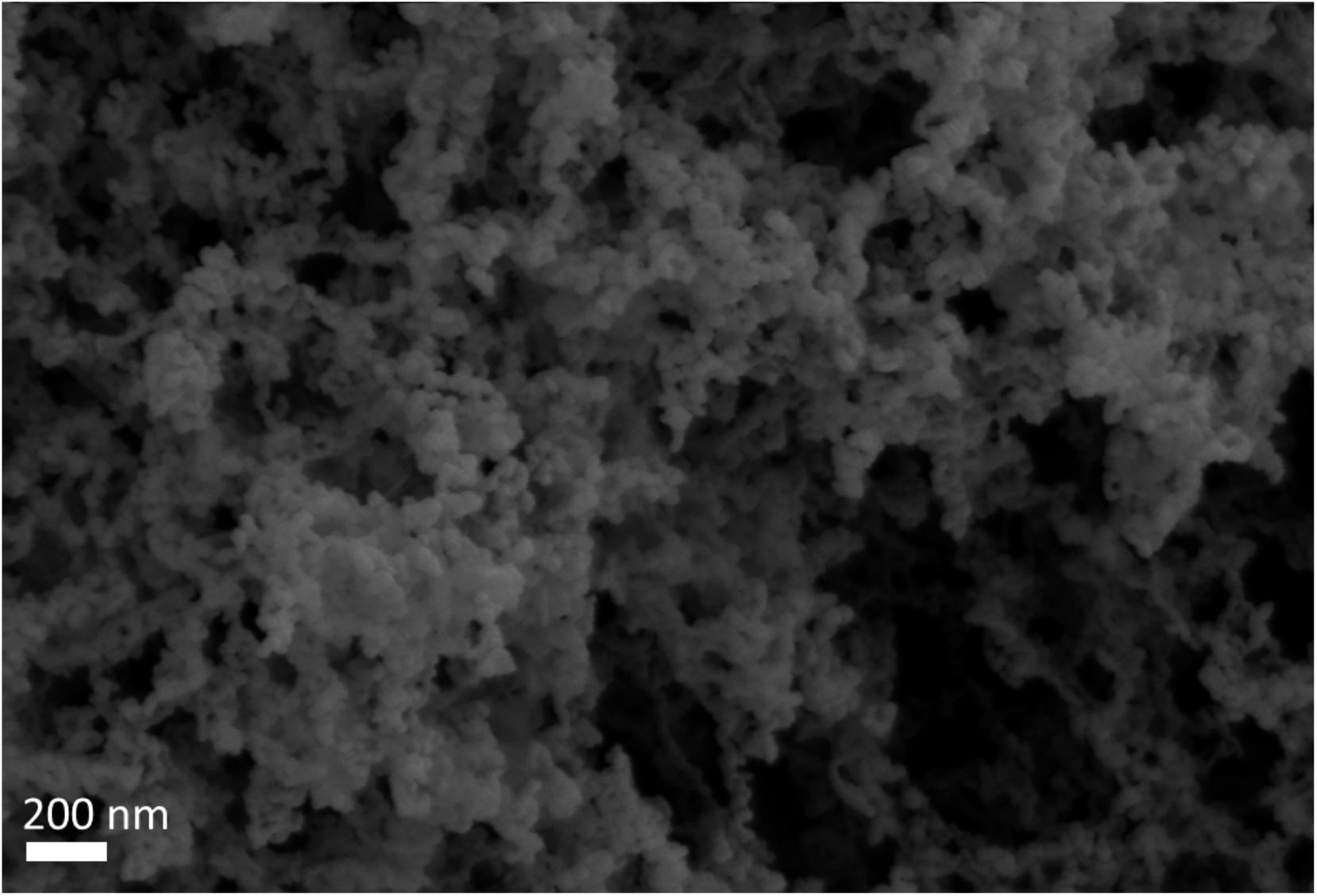
SEM picture of niclosamide inhalation powder. The bar shows a size of 200 nm.

### Multi-dose administration of niclosamide in rats confirmed that it is slowly cleared from the lungs with mild pulmonary toxicity

We performed a multi-dose pharmacokinetic study on inhaled niclosamide for the purpose of obtaining information on the potential toxicology of inhaled niclosamide after administration of a relatively high dose (~827.2 μg/Kg of niclosamide contained in the TFF inhalation powder described in Table 1 for rats). The steady-state plasma drug concentration after three days of daily dosing was chosen to assess the accumulation and absorption of niclosamide. The results of the plasma and lung concentrations of niclosamide are provided in Table 3. The amount of niclosamide in the lungs at 24 hours following administration of the last dose was undetectable in two out of five rats and less than 1 ng/mL in all five of the tested rats. Concentrations in the ng/mL range were still detected in the rat plasma samples from four out of five rats. All rats survived between administration and sample collection.

**Table 3.**
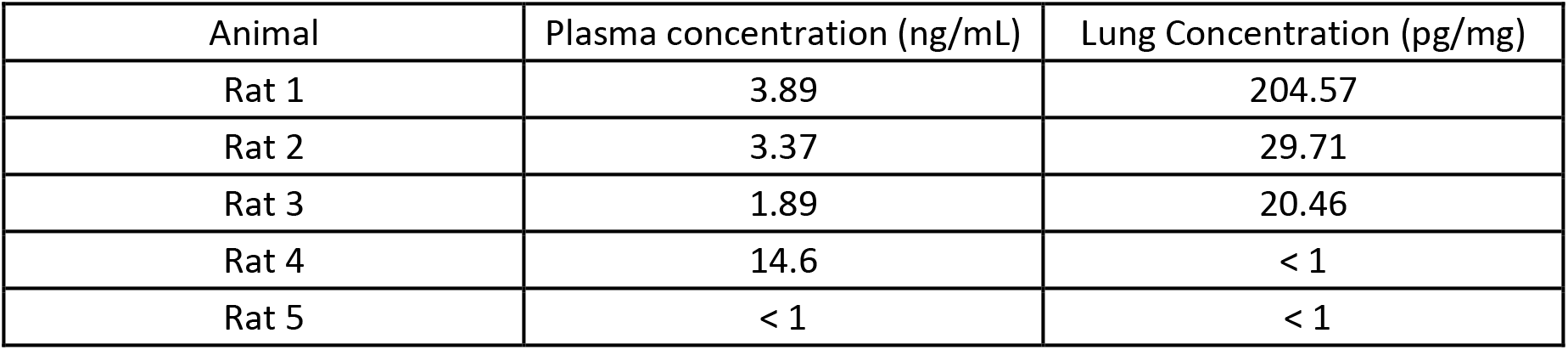
Concentration of niclosamide in the plasma and lungs of Sprague-Dawley rats after three days of daily dosing of niclosamide inhalation powder (200 μg of niclosamide daily per animal, ~827.22 μg/Kg). The limit of quantification in plasma samples was 1 ng/mL and 1 pg/mg in lung tissue.

The histopathological examination of the liver, kidney, and spleen from the treated rats appeared normal. However, examination of the lung tissue indicated mild signs of inflammation after daily dosing for three days, which is likely related to the relatively high doses of niclosamide inhalation powder that were administered (niclosamide, leucine, and mannitol). In all animals, the air space and interstitial space were found to be normal. The examination also indicated a few to moderate eosinophils in the lung tissue with rare to scattered polymorphonuclear (PMN) cells observed. This was accompanied by some bronchitis in most of the rats. The specific findings in each animal are presented in Table 4. A representative image from Rat 4 is provided in Figure 5, depicting normal alveoli (Figure 4A) and some bronchial inflammation (Figure 4B) with mixed granulocyte and eosinophils present. In one animal, Rat 3, all findings were normal.

**Table 4.** Histopathology observations in each of the treated Sprague-Dawley Rats after three days of dosing niclosamide inhalation powder (200 μg/niclosamide administered daily per animal).

| Animal | Histopathological Findings |
| --- | --- |
| Rat 1 | Air space and interstitial space normal; normal kidney, liver, and spleen; bronchovascular bundle with moderate eosinophils with admixed PMNs. Large airways > small airways with mild-moderate bronchitis. |
| Rat 2 | Air space and interstitial space normal; normal kidney, liver, and spleen; parabronchial eosinophils only in the large airways; otherwise predominantly normal. |
| Rat 3 | Air space and interstitial space normal; normal kidney, liver, and spleen; lungs normal. |
| Rat 4 | Air space and interstitial space normal; normal kidney, liver, and spleen; some eosinophilic bronchitis; scattered PMN with patches of bronchitis among some normal airways. |
| Rat 5 | Air space and interstitial space normal; normal kidney, liver, and spleen; moderate eosinophilic bronchitis with few PMNs. |

**Figure 4.**
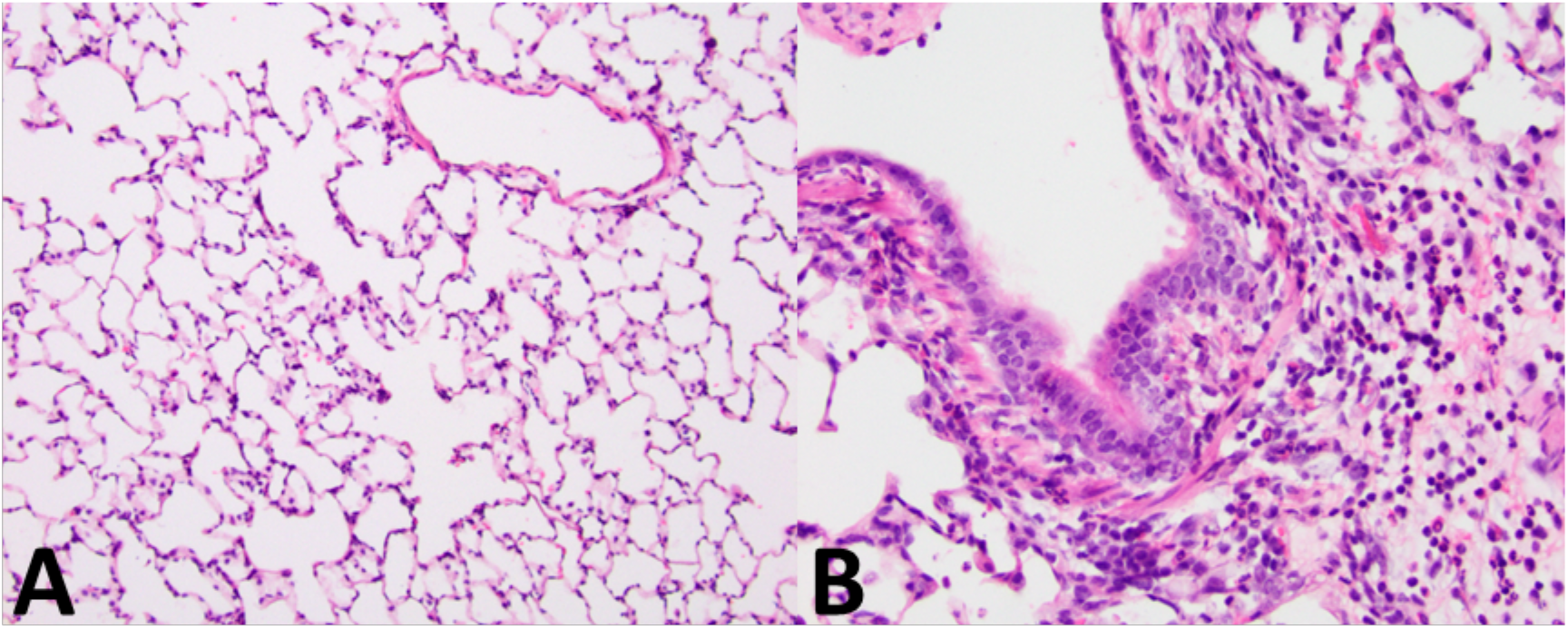
(A) Histopathological examination of lung tissue showing normal alveolar region at a magnification of 100x and (B) bronchial inflammation of a 1 mm airway section in Rat 4 after three days of dosing niclosamide inhalation powder (200 μg of niclosamide per animal).

**Figure 5.**
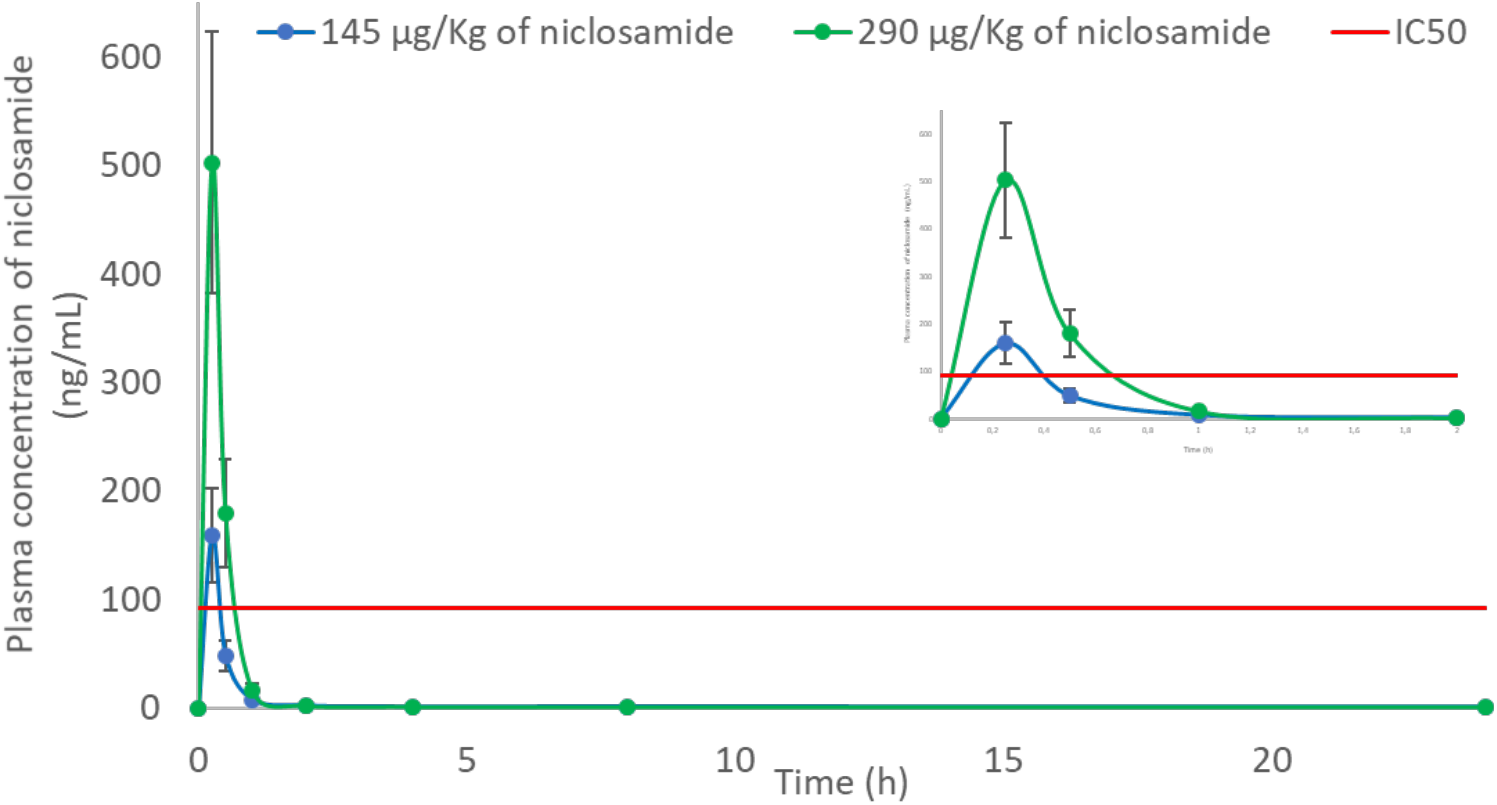
Plasma concentration profile of niclosamide after the administration of niclosamide inhalation powder in Syrian hamsters (n=5 for each point). The groups received 8.7 (blue) and 17.4 (green) mg/Kg of niclosamide inhalation powder, containing a dose of 145 and 290 μg/Kg of niclosamide, respectively. The inset picture shows the profile within the first 2 h. Niclosamide IC_50_ for SARS CoV 2 is 0.28 μM (91.56 ng/mL) (red)[8].

Chang *et al*. (2006) reported a T_1/2_ of 6.0 ± 0.8 h for solubilized niclosamide after oral administration [35]. Based on the measured niclosamide concentrations in Sprague-Dawley rat plasma from Table 4 after 24 h of administration (~ 4 x T_1/2_), there appears to be a prolongation in its apparent half-life as it was possible to detect and quantify niclosamide in rat plasma. Moreover, our group recently reported that the oral administration of niclosamide suspension had a T_1/2_ of approximately 1.0 ± 0.3 h in Wistar rats [36]. So, the results of niclosamide inhalation powder and those concentrations above the limit of quantification support the idea of once-daily administration. This may be attributed to an extended residence time in the lung resulting from the avoidance of the extensive first-pass effect that niclosamide undergoes after oral administration [37]. Despite the fact that some drug remains in the plasma, it is reasonable that niclosamide does not accumulate in the lungs following three consecutive days of daily dosing as supported by the low concentrations measured in the lung tissue (Table 4).

Costabile *et al*. 2015 performed an acute lung toxicity study of niclosamide nanosuspensions (containing mannitol) in Wistar rats (200 and 220 g). They did not observe acute toxicity after intratracheal administration of the niclosamide nanosuspension at doses ranging from 10 to 100 μg per animal [15]. In our work, the Sprague-Dawley rats (252.4 – 279.5 g) were administered 200 μg/rat daily for three days, and some signs of mild to moderate inflammation were observed. The mild to moderate inflammation of the bronchioles along with the presence of eosinophils at this dose is a promising sign and currently unclear if these observations result from niclosamide or the mannitol excipient in the formulation, which is reported to cause bronchospasm when administered at high milligram doses in humans [38].

### Niclosamide inhalation powder sustains lung concentrations above the IC_50_ and estimated IC_90_ for 24 hours in the Syrian Golden Hamster model

The *in vivo* pharmacokinetics and toxicology study of niclosamide inhalation powder in rats at the dose of ~827.2 μg/Kg confirmed the feasibility of the formulation to be translated to a COVID-19 infection model. The Syrian golden hamster model is relevant for SARS-CoV-2 infection, and a complete pharmacokinetic analysis is required in order to evaluate the proper dosing for further efficacy studies [39,40]. The hamsters were administered niclosamide inhalation powder prepared at a lower niclosamide concentration in order to have a dose of 145 and 290 μg/kg niclosamide. As can be seen in Figure 5, there was fast absorption of niclosamide that was rapidly cleared to steady low levels; in general, this is considered an advantage of inhaled drug products in terms of safety and reduction of systemic side effects [41]. A non-compartmental analysis of the plasma data was performed; the calculated pharmacokinetic parameters are shown in Table 5. This analysis showed dose proportionality between the doses as evidenced by the AUC/dose of both administered doses are similar (8.6 and 8.4), and clearance was in the range of ~116-119 mL/h. All hamsters survived between administration and sample collection.

**Table 5.** Plasma in vivo pharmacokinetic parameters of niclosamide inhalation powder in two groups of Syrian hamsters. The groups received 8.7 and 17.4 mg/Kg of niclosamide inhalation powder (described in Table 1 for Syrian golden hamsters), containing a dose of 145 and 290 μg/Kg of niclosamide, respectively.

| Pharmacokinetic parameters | Niclosamide dose of 145 $\mu\text{g/Kg}$ | Niclosamide dose of 290 $\mu\text{g/Kg}$ |
| --- | --- | --- |
| $T_{1/2}$ (h) | 53.5 | 41.6 |
| $T_{\text{max}}$ (h) | 0.3 | 0.3 |
| $C_{\text{max}}$ (ng/mL) | 159.0 | 590.3 |
| $AUC_{\text{last}}$ (h*ng/mL) | 83.8 | 237.1 |
| $AUC_{\text{Inf}}$ (h*ng/mL) | 137.9 | 257.1 |
| AUC_infinity_dose | 8.6 | 8.4 |
| $AUC_{\% \text{Extrap}}$ | 39.2 | 7.8 |
| CL (mL/h) | 116.8 | 119.4 |
| $MRT_{(h)}$ | 41.5 | 7.3 |

Furthermore, in the lung tissue (Figure 6), a similar trend was observed, and the steady-state level of niclosamide exceeded the IC_50_ for at least 24 h. Unfortunately, the lung concentration profile was not possible to determine using the non-compartmental analysis model, probably due to the inability to observe the elimination phase of niclosamide in lung tissue between 4 and 24 h. Therefore the only parameters reported are C_max_, AUC _0-24h_, and T_max_ (Table 6).

**Figure 6.**
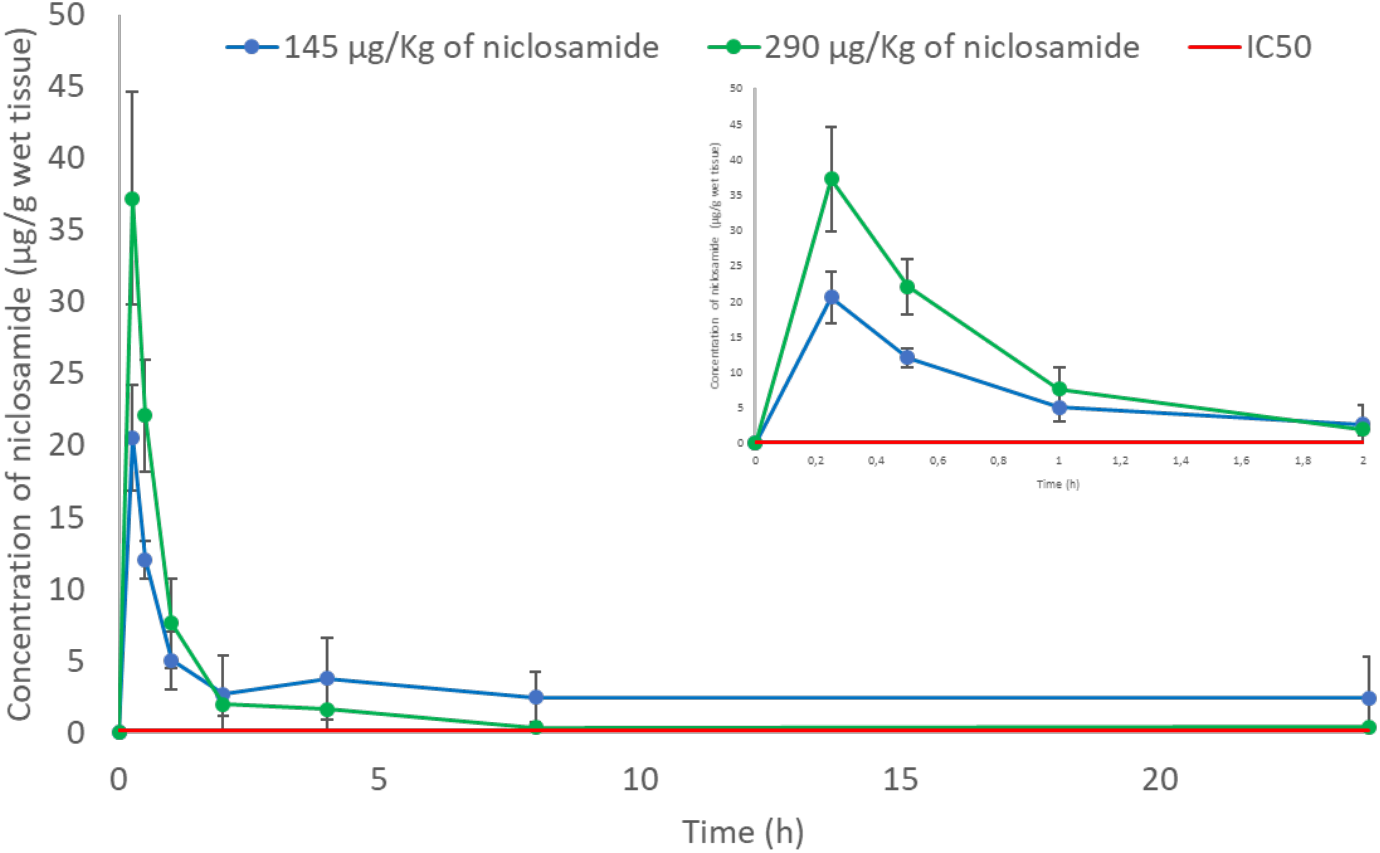
Lung concentration profile of niclosamide after the administration of niclosamide inhalation powder to Syrian golden hamsters (n=5). The groups received 8.7 (blue) and 17.4 (green) mg/Kg of niclosamide inhalation powder, containing a dose of 145 and 290 μg/Kg of niclosamide, respectively. The inset picture shows the profile within the first 2 h. Estimated niclosamide IC_50_ for SARS CoV 2 is 0.09156 μg/g of wet tissue (red), it was assumed that the wet tissue has a density of 1 g/mL.

**Table 6.**
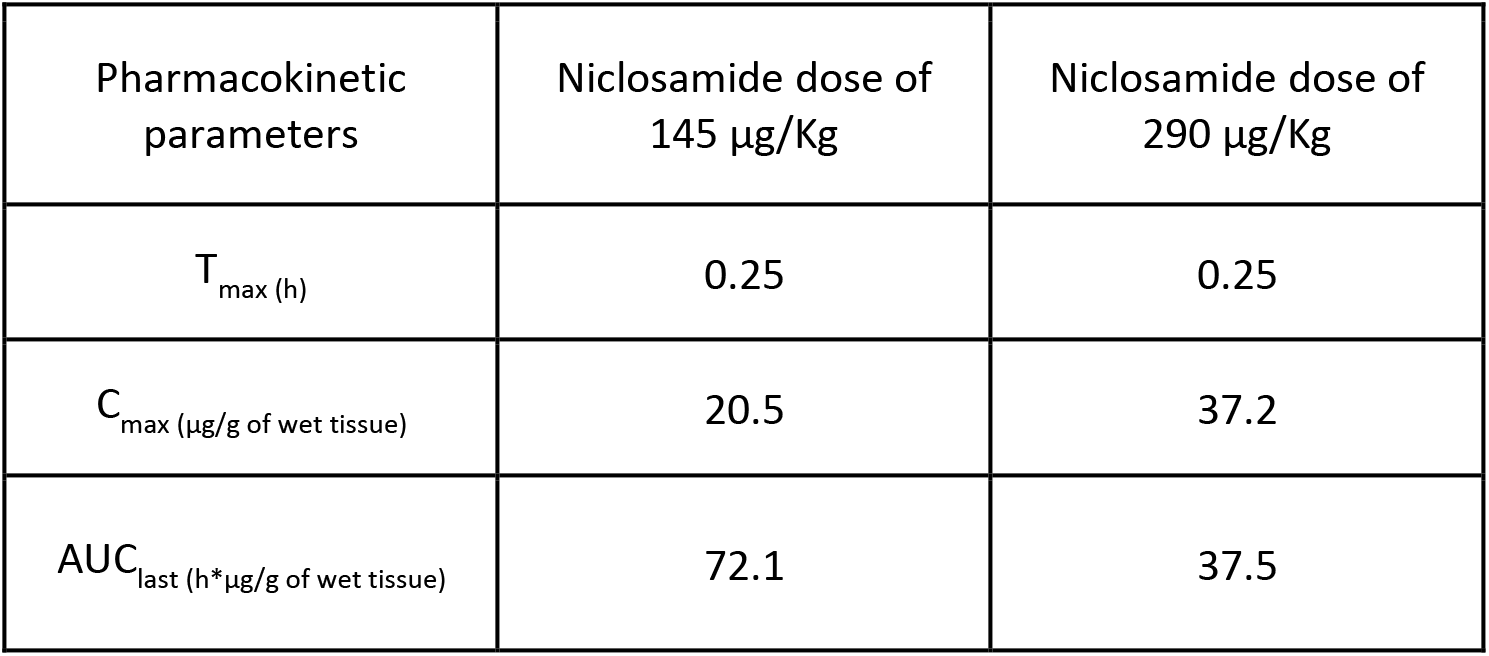
Lung in vivo pharmacokinetic parameters of niclosamide inhalation powder in two groups of Syrian hamsters (described in Table 1 for Syrian golden hamsters). The groups received 8.7 and 17.4 mg/Kg of niclosamide inhalation powder, containing a dose of 145 and 290 μg/Kg of niclosamide, respectively.

| Pharmacokinetic parameters | Niclosamide dose of 145 μg/Kg | Niclosamide dose of 290 μg/Kg |
| --- | --- | --- |
| $T_{max}$ (h) | 0.25 | 0.25 |
| $C_{max}$ (μg/g of wet tissue) | 20.5 | 37.2 |
| $AUC_{last}$ (h*μg/g of wet tissue) | 72.1 | 37.5 |

There was a notable difference in the plateau concentration achieved by the two different administered doses in terms of the concentrations reached in the lungs after 2 h. The lower administered dose had a higher AUC_0-24 h_ without achieving a higher C_max_. We hypothesize that the administration of a higher dose of powder caused a change in aerosolization properties from the dry powder insufflator used to administer the dose of powder to the hamsters. Molina *et al*. (2016) compared different methods of pulmonary administration of nanoparticles, and they found that with dry powder insufflation, the nanoparticles were cleared faster because the powder was deposited in the larger airways and trachea with higher variability among the animals (i.e., 188%). Moreover, they detected higher amounts of their formulation in the gastrointestinal tract, suggesting mucociliary clearance from the upper respiratory tract. They speculated that the deposition in the trachea was related to issues in the synchronization between the insufflation and the inhalation cycle of the animals and not due to properties of the drug [42]. The work of Duret *et al*. (2012) confirms that the DP-4M^®^ insufflator (used in this study) is reliable for delivering different drug doses of the same powder in mice, but unfortunately, in their work, the animals were immediately sacrificed to measure drug deposition [43]. So, no relationship between deposition in different regions of the lungs and their effect on pharmacokinetics was established. Moreover, Hoppentocht *et al*. (2014) proved that the insufflator is not consistent or efficient in deaggregating powders using 200 μL of air pulsed at different doses [44].

Based on these reports of using dry powder insufflators in animal studies, the differences observed between the dose amounts administered in our study and the expected lung pharmacokinetics is likely related to the efficiency of deaggregation of the two-dose amounts delivered from the dry powder insufflator and not related to niclosamide itself. This is because changes in powder deposition caused by the insufflator can lead to different emitted particle sizes from the device and subsequent different dissolution/solubility of the emitted particles, mechanisms of clearance, and drug permeation [45]. The C_max_ of niclosamide following dosing by insufflation to the lungs is nearly doubled as the dose was doubled, which agrees with the studies that showed that the insufflator is reliable for delivering the loaded dose.[43] However, it is known that particles deposited in the trachea and larger airways are cleared faster. [46] This is supported by the plasma AUC_0-24 h_ that was more than double when the dose was doubled. So, a greater amount of niclosamide from the powder formulation was absorbed and cleared by the plasma instead of being retained in the lung tissue (Table 5). Huang *et al*. (2018) used the coefficient C_lung_/C_plasma_ to describe the biodistribution of an itraconazole dry powder inhalation product in a pharmacokinetic study [47]. Due to the differences in concentrations from Huang *et al*., we used the reciprocal coefficient, C_plasma_/C_lung_ to describe the distribution between plasma and lungs. As can be seen in Figure 7, niclosamide inhalation powder behaves differently after being administered at two different dose amounts to the hamsters; a greater amount of the drug was detected in plasma than in lungs after increasing the administered dose. This supports our argument that at the higher dose amounts loaded and delivered from the dry powder insufflator, the insufflated powder was likely deposited in the larger airways, which can impact the pharmacokinetics of the administered dry powder by increasing the clearance of particles [48].

**Figure 7.**
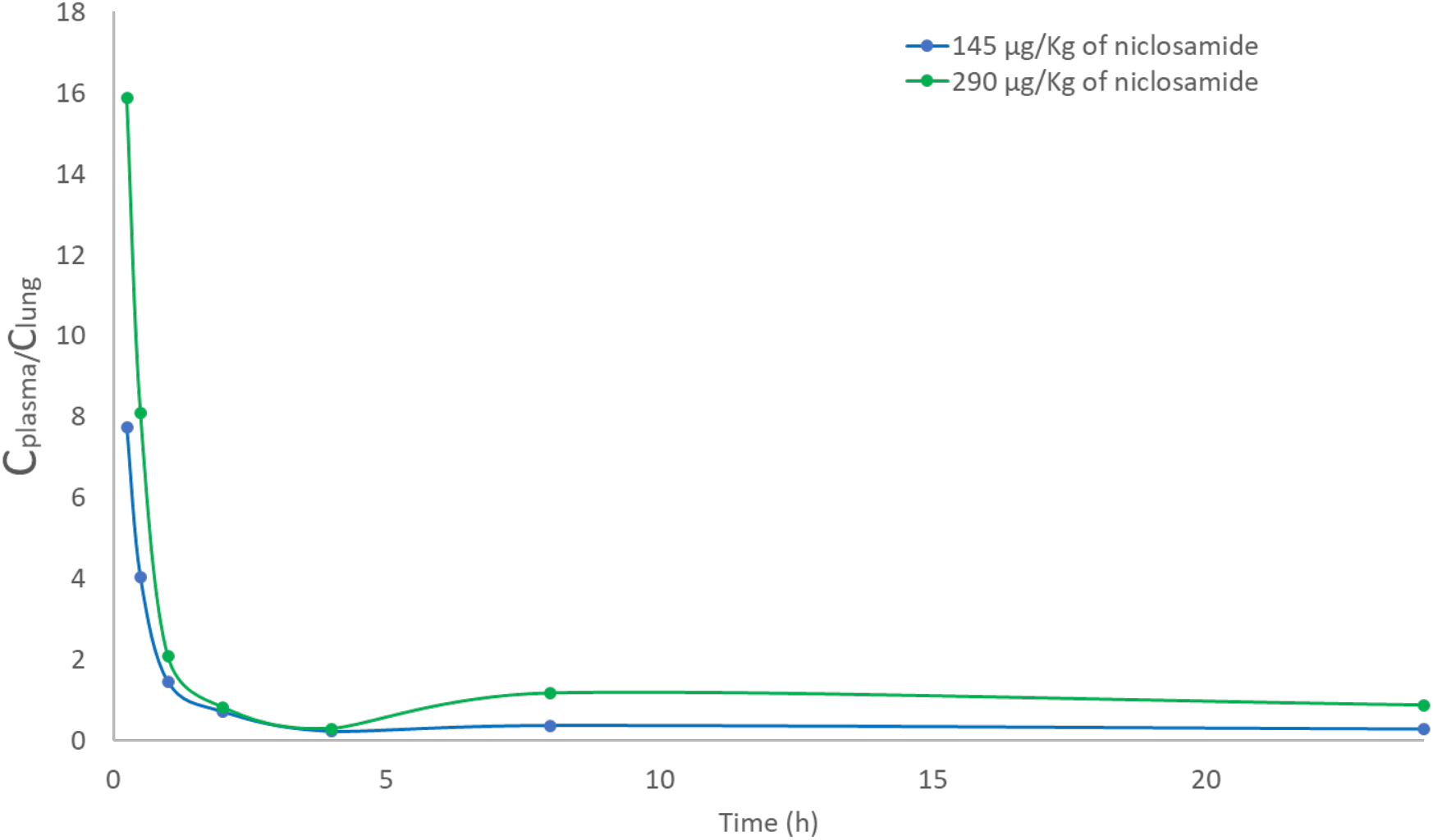
Coefficient of niclosamide concentration in plasma and lung over time (C_plasma_/C_lungs_). The groups received 8.7 (blue) and 17.4 (green) mg/Kg of niclosamide inhalation powder, containing a dose of 145 and 290 μg/Kg of niclosamide, respectively.

To the best of our knowledge, the present work is the first describing the pharmacokinetics of niclosamide following delivery to the lungs. It is relevant and informative to compare our results to pharmacokinetic studies reported that delivered niclosamide by other routes of administration and animal model. Ye *et al*. (2015) administered niclosamide intravenously (2 mg/Kg) through the jugular vein to Sprague–Dawley rats using a mixture of solvents and measured the drug’s biodistribution in the liver, heart, spleen, lungs, kidneys, and intestines. The concentration in the lungs and heart were consistently the lowest based on intravenous delivery [49]. Tao *et al*. (2014) reported the concentrations of niclosamide in male C57BL/6J mice after oral administration of niclosamide ethanolamine at 40 mg/Kg. In their work, the lung concentrations following oral administration were decreased from 87.2 ± 70, 23.8 ± 13, and 13.3± 5 ng/g at 2, 4, and 8 h, respectively. Interestingly, the ethanolamine salt form of niclosamide mainly distributed to the liver, and higher levels of metabolites were detected in the kidneys. Therefore, neither intravenously administered niclosamide nor orally administered niclosamide ethanolamine provided therapeutically relevant lung levels of niclosamide, thus emphasizing the importance of the inhaled route of administration of niclosamide directly to the lungs for lung infections like SARS-CoV-2.

By considering the differences in animal models, Figure 6 shows that it is possible to achieve lung concentrations mostly in the μg/g scale and above the IC_50_ using lower doses just in the range of μg/Kg for at least 24 h because the drug was targeted directly into the lungs. The development of niclosamide inhalation powder avoids first-pass metabolism and reduces the exposure of the drug to being metabolized by hydroxylation and glucuronidation [50].

Finally, in further considering administration to infected patients, the inhaled niclosamide powder should also achieve concentrations above the IC_90_. In our hamster studies, we administered a TFF inhaled powder formulation containing a reduced amount of niclosamide (i.e., 1.66% (w/w) dosed at 145 and 290 μg/Kg to hamsters compared to 22% (w/w) dosed at 827.22 μg/Kg to rats). Based on the reported inhibitory effect of niclosamide on SARS-CoV-2 and an IC_50_ of 91.56 ng/mL (0.28 μM) following the work of Jeon *et al*. (2020), Arshad *et al*. (2020) estimated the IC_90_ of 153.7 ng/mL (the paper considered IC_50_ = EC_50_). To be truly useful as a therapeutic against SARS-CoV-2, the niclosamide inhalation powder should achieve a concentration higher than 0.154 μg/g, assuming that the wet tissue has a density of 1 g/mL (assumed to be equal to the density of water). We exceeded this concentration in lung tissue using both administered doses by more than 2-fold over the 24 h period (the lower dose was ~15-fold) as seen in Table 6. Further supporting our analysis, comparisons with the report of Gassen *et al*. (2020) is helpful. The paper reports an IC_50_ of 55.6 ng/mL (0.17 μM) [51]. Therefore, niclosamide inhalation powder is also above this concentration. Additionally, the toxicology studies support that niclosamide inhalation powder has a sufficient safety margin for dose adjustment, thus allowing for a once per day inhalation therapy.

## Conclusion

Using thin-film freezing, we report the administration of niclosamide inhalation powder to rats and Syrian golden hamsters. The TFF powders have remarkable aerosol properties for targeting the lungs, and especially, the lower respiratory tract. The niclosamide inhalation powder was demonstrated to be safe after pulmonary administration in the rat model, and the pharmacokinetics study in Syrian golden hamsters, an important animal model for SARS-CoV-2 infection, showed that the drug remains in the lungs for at least 24 h at concentrations greater than the reported IC_50_ and IC_90_ for SARS-CoV-2 after just a single inhaled dose administered by inhalation.

## Funding

This research was funded by TFF Pharmaceuticals, Inc. through a sponsored research agreement with the University of Texas at Austin. Warnken is partially supported by this sponsored research agreement with TFF Pharmaceuticals Inc. Miguel O. Jara acknowledges the funding support from the Equal Opportunities Fulbright—CONICYT Scholarship 56170009.

## Conflicts of Interest

Jara, Warnken, Moon, Sahakijpiarn and Williams are co-inventors on IP related to this paper. The University of Texas System has licensed this IP to TFF Pharmaceuticals, Inc. Moon and Sahakijpijarn acknowledge consulting for TFF Pharmaceuticals, Inc. Williams owns equity in TFF Pharmaceuticals, Inc. Peters is a consultant to TFF Pharmaceuticals, Inc. (uncompensated).

## Supplemental Data File for Jara et al, Inhaled Niclosamide in Rats and Hamsters

**Table S1.** The triple quadrupole MS was operated in electrospray positive ion detection in MRM mode detecting the following transitions, where Q indicates the quantifying MRM transition and C indicates the confirming MRM transition. The instrument was set up to divert the first 2.7 minute to waste. The dwell time for each MRM transition was 100 msec.

| Compound | parent m/z | product m/z | fragmentor (V) | Collision energy (V) |
| --- | --- | --- | --- | --- |
| Niclosamide (Q) | 324.9 | 171 | 120 | 25 |
| Niclosamide (C) | 324.9 | 269 | 120 | 15 |
| <sup>13</sup>C<sub>6</sub> Niclosamide (IS-C) | 330.9 | 177 | 120 | 25 |
| <sup>13</sup>C<sub>6</sub> Niclosamide (IS-Q) | 330.9 | 295.1 | 120 | 15 |

